# Rapid onset fibrotic remodeling and ventricular dysfunction induced by phenylephrine involve targeted reprogramming of myocyte and fibroblast transcriptomes

**DOI:** 10.1101/2024.10.11.617933

**Authors:** Todd H. Kimball, Tatiana Gromova, Natalie D. Gehred, Douglas J. Chapski, Ke Wang, Marmar Vaseghi, Matthew A. Fischer, David J. Lefer, Thomas M. Vondriska

## Abstract

Catecholamine dysregulation is a common feature of multiple acute and chronic cardiac conditions, including heart failure. To investigate the role of altered α-adrenergic stimulation on cardiac function, we developed a short-term exposure model, administering phenylephrine subcutaneously to mice for one week. Compared to vehicle-injected controls, phenylephrine-treated animals exhibited increased ejection fraction, decreased chamber size, diastolic dysfunction and ventricular hypertrophy in the absence of hypertension. Remarkably, these animals developed extensive fibrotic remodeling of the tissue that plateaued at 24 hours and myocyte hypertrophy localized to regions of fibrotic deposition after 3 days of treatment. Transcriptome analyses of purified myocyte and fibroblast populations from these hearts revealed an unexpected role for myocytes in the production of extracellular matrix. Comparison with other models of cardiac stress, including pressure overload hypertrophy and cytokine activation of fibroblasts, identified stimulus-specific transcriptional circuits associated with cardiac pathology. Given the rapid, robust fibrotic response that preceded myocyte hypertrophy, intercellular communication analyses were conducted to investigate fibroblast to myocyte signaling, identifying potential crosstalk between these cells. These studies thoroughly describe and phenotypically characterize a new model of short-term catecholamine stress and provide an atlas of transcriptional remodeling in myocytes and fibroblasts.

**Graphical Abstract:** 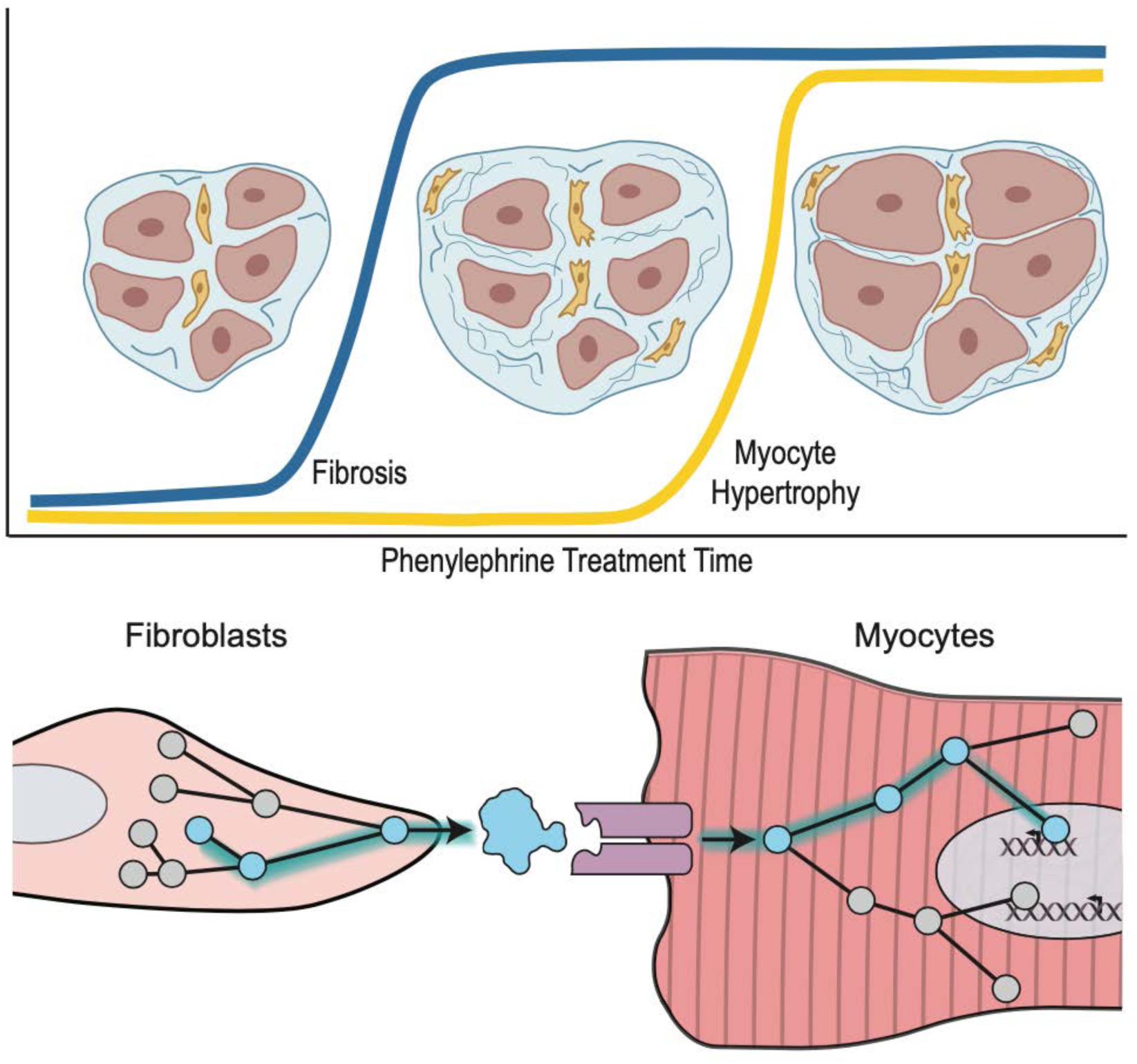

## Introduction

Heart failure is a chronic syndrome that develops in response to varied stressors, including hypertension, excessive adrenergic stimulation, pressure overload, inflammation and other stimuli. Pathological hypertrophic remodeling, phenotypically characterized by a thickening of the left ventricular wall and septum and interstitial fibrosis, stiffens the heart and induces cell-specific responses to the altered environment. Cardiomyocytes enlarge to increase contractility, and fibroblasts exit a quiescent state to adapt a proliferative phenotype or to undergo myofibroblast differentiation and extracellular matrix secretion. Acute adrenergic signaling induces cardiomyocyte protein synthesis, as sarcomeric proteins organize in parallel to increase cardiomyocyte width, and fibroblast proliferation. These two processes drive concentric hypertrophy, which increases left ventricular wall thickness and tissue mass, marking a compensatory phase of the cardiac stress response. Heart failure with reduced ejection fraction is a syndrome in which low cardiac output and tissue perfusion leads to a state of elevated catecholamine release, which can precipitate lethal arrhythmias ^1^. In this case, a weakened heart muscle, often compromised by infarction, fibrotic scarring and metabolic dysfunction, cannot respond to adrenergic stimulation by increasing contractility *and* the subject’s impaired ambulatory ability prevents systemic skeletal muscle use. While the gene expression of cardiac cell types under stress has been characterized, the cross talk between cell types are still being uncovered.

Catecholamines have important actions on the heart that manifest during acute and chronic pathophysiological conditions. α and β receptor agonists can induce hypertrophy by direct actions on myocytes^2^ and these compounds drive fibrotic responses in the myocardium ^3^ through their actions on fibroblasts, myocytes and potentially other cells. Acute administration of phenylephrine, an α-adrenergic agonist, induces gene activation distinct from β-adrenergic stimulation ^4^, concomitant with myocyte hypertrophy *in vivo* ^5^ as well as in isolated cells ^6^. During injury such as ischemia or the chronic pressure overload associated with hypertension, it has long been appreciated that multiple cell types in the heart, including myocytes, fibroblasts, vascular cells and immune cells contribute to the short- and long-term healing processes. More recent studies have employed single cell or single nucleus RNA sequencing to address this question, measuring changes in transcription across the totality of experimentally accessible cells in the tissue ^7^. The advantage of these studies is that many cells are individually sampled across a time course of injury and recovery, such as following myocardial infarction^8^ or pressure overload ^9^. A limitation of the single cell or single nucleus RNA sequencing approach, however, is that the experimenter determines *a priori* the identity of a revealed cell population based on the markers chosen to demarcate it. In other words: one can sample the tissue in an unbiased manner, but the data analysis proceeds according to known cell marker genes and in the absence, in most cases, of orthogonal identifying phenotypes of the cells, such as morphology.

We examined the heart’s response to stress in the early stages of disease progression to elucidate the network of communication between cell types that gets hijacked to propagate pathologic signaling. We sought to investigate the direct impact of catecholamine administration on cardiac function and the commensurate short-term transcriptome reprogramming that occurs in response to this stress. While longer term administration of catecholamines like isoproterenol is better understood ^10, 11^, the early, rapid response of myocyte and fibroblast transcriptomes to catecholamine stimulation is less well characterized and the effects of phenylephrine on acute phenotypes and gene expression is understudied. Human cardiac pathologies including hypertrophic obstructive cardiomyopathy ^12^, Takotsubu syndrome ^13^, myocardial ischemia ^14^ and longer term heart failure ^15^ are known to involve aberrant catecholamine signaling. In the case of heart failure, β-adrenergic receptor stimulation is well known to promote the incidence and progression of this syndrome ^15^, yet α-adrenergic receptor signaling exerts an interestingly nuanced adaptive response, promoting protection from ischemia and physiologic hypertrophy ^16^. Stimulation of α-adrenergic receptors with phenylephrine in humans was shown to have a positive ionotropic effect ^17^. Recent studies have examined the ability of short-term phenylephrine to induce cardiac muscle stiffening and adaptation in mice ^18^.

In the present study, mice treated with phenylephrine for 7 days developed myocyte and ventricular hypertrophy, elevated ejection fraction associated with a reduced chamber volume, and diastolic dysfunction, in the absence of sustained hypertension. Detailed examination of tissue fibrosis revealed a peak after one day and lack of further fibrotic progression. Areas of enhanced matrix deposition co-localized with regions of myocyte hypertrophy. To investigate the transcriptional reprogramming induced by phenylephrine, cardiac myocytes and fibroblasts were separately isolated using established protocols^19, 20^ that enrich for these cell types as determined by morphological and functional analyses. We interrogated these transcriptomes in comparison to reprogramming of myocytes and fibroblasts in response to other pathologic stimuli, including pressure overload and cytokine stimulation. Our findings provide an atlas of cardiac transcription in response to acute catecholamine stimulation and reveal both myocytes and fibroblasts to be key contributors to extracellular matrix deposition in the setting of cardiac hypertrophy and fibrosis.

## Results

Phenylephrine (or PBS control) was administered subcutaneously every other day over 7 days to adult, male, C57Bl/6J mice at which point cardiac function was assessed by echocardiography (tissues were harvested on day 8 for histology and RNA-seq; Figure 1A). Phenylephrine induced thickening of the ventricular anterior and posterior wall, in turn decreasing left ventricular chamber size while increasing ejection fraction (Figure 1B, 1C). Fractional shortening was also increased, whereas global longitudinal strain was decreased in the phenylephrine treated animals, concomitant with cardiac hypertrophy as measured by heart weight to tibia length or heart weight to body weight ratios (Supplemental Figure 1 and Figure 1E). Invasive hemodynamics showed slight blood pressure decrease in phenylephrine treated animals compared to PBS (Supplemental Figure 1). To examine cardiac relaxation, we measured E/A ratio, which reflects ventricular stiffness, and found it elevated (Figure 1D). Phenylephrine treated mice also exhibited decreased e’ (Supplemental Figure 1) and elevated E/e’ ratio (Figure 1D), consistent with diastolic dysfunction^21^. In addition, mean LVEDP measured by cardiac catheterization was higher in phenylephrine treated mice, but the difference was not statistically significant (Supplemental Figure 1). Together, these data demonstrate ventricular stiffening and rapid adaptive growth leading to impaired cardiac output in response to short term α-adrenergic stimulation.

**Figure 1.**
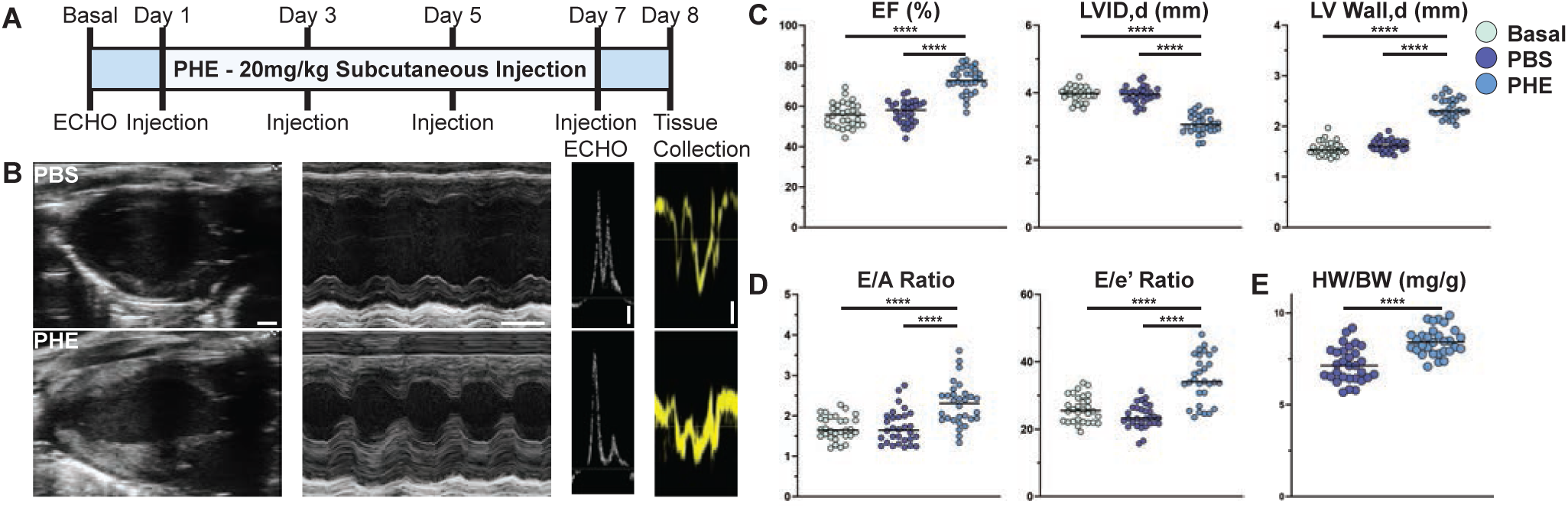
Acute cardiac dysfunction following α_1_-adrenegic stimulation. **A.** Timeline of phenylephrine administration (PHE, 20 mg/kg) or Vehicle (PBS) as subcutaneously injected every other day for 7 days, with echocardiographic (ECHO) parameters measured on Day 0 (basal) and Day 7, tissue was harvested on Day 8. **B.** Representative echocardiographic images from PBS or PHE treated mice on Day 7 to observe systolic and diastolic phenotypes. From left to right, Long Axis B Mode (scale bar = 1 mm), Short Axis M Mode (scale bar = 100 msec), Pulse Wave Doppler (scale = 100 mm/s), and Tissue Doppler (scale = −10 mm/s). **C.** Quantification of systolic function at Basal (green), PBS (violet) and PHE (blue). PHE increased ejection fraction (EF(%)) and left ventricular wall thickness during diastole (LV Wall,d), while decreasing left ventricular chamber diameter (LVID,d). **D.** Quantification of diastolic function. PHE increased E/A and -E/e’ ratios, indicative of diastolic dysfunction (n= 30 mice/group, ****p<0.0001, One-Way ANOVA and Tukey’s multiple comparisons test for systolic and diastolic measurements). **E.** PHE treated group had a significantly increased heart weight/body weight ratio (HW/BW) (n= 30 mice/group, ****p<0.0001, Welch’s t test).

We next characterized the histopathologic manifestations of this phenylephrine-induced cardiac dysfunction. Phenylephrine treatment caused narrowing of the left ventricular chamber and thickening of the walls as evident in longitudinal sections of hearts stained with picrosirius red, which at higher magnification clearly demonstrate significant accumulation of interstitial fibrosis (Figure 2A and 2C). Wheat germ agglutinin staining of cell membranes was used to visualize cardiomyocyte morphology and quantify cell size changes, revealing hypertrophy at the myocyte level (Figure 2B and 2D). Different regions of the ventricle experience substantially different physical forces during contraction and host different populations of adrenergic receptors ^22^. To more precisely investigate the anatomical differences in myocyte hypertrophy and the deposition of fibrosis, we examined pathologic changes in the interventricular septum, left ventricular free wall and apex. Moreover, we sought to quantify the interplay between myocyte growth and fibroblast activation by separately examining myocyte size in fibrotic versus non-fibrotic regions of these left ventricular locations. In the ventricular free wall and the apex, hypertrophy occurred only in areas with significant fibrosis, whereas in the interventricular septum, fibrotic and non-fibrotic areas were populated with hypertrophied myocytes (Figure 2E). To investigate the temporal dynamics of this phenomenon, we measured fibrosis and myocyte size in separate cohorts of mice sacrificed at various time points after initial phenylephrine administration. The fibrotic response was rapid, appearing as early as one day and plateauing out to a week, whereas the increase in myocyte size was more gradual, plateauing around day 3 (Figure 2F). These observations demonstrate that fibrosis precedes hypertrophy in response to phenylephrine and suggest that extracellular matrix deposition (or other secretory behavior) by fibroblasts may drive the hypertrophic cardiomyocyte response.

**Figure 2.**
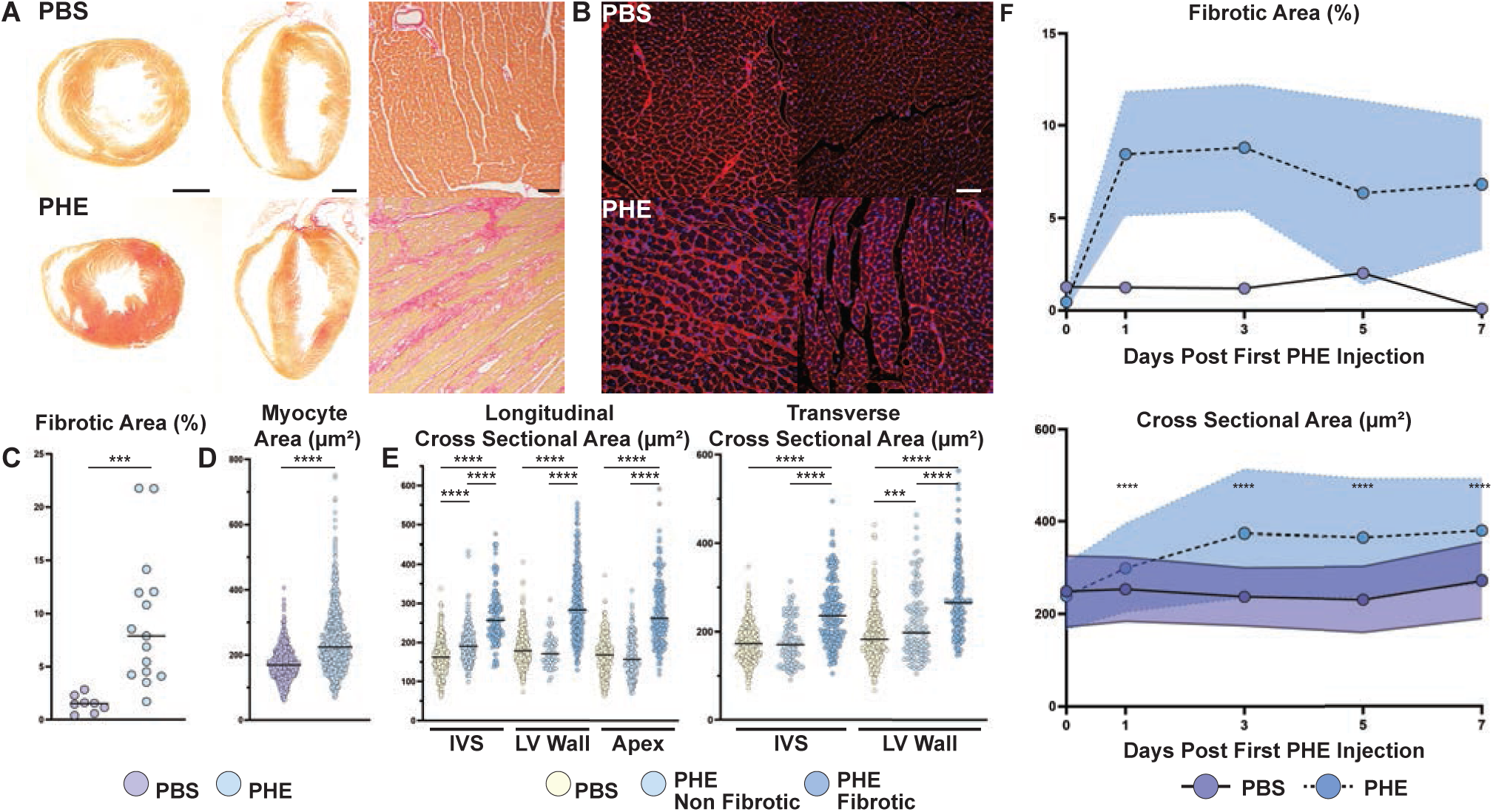
Phenylephrine induced cardiac tissue and cellular remodeling. **A.** Picrosirius staining of cardiac tissue sections. Transverse (far left, scale = 1mm)) and Longitudinal (middle, scale = 1mm) sections of PHE show thickened interventricular septum, LV wall, and apex (longitudinal only). PHE increased fibrotic deposition (right, scale = 50 µm) (n= 4 PBS, n=7 PHE). **B.** Fluorescent microscope imaging of wheat germ agglutin (WGA) stained cardiac tissue, showing increased density after PHE (scale = 50 µm). **C.** Whole tissue fibrotic area quantification (analysis with FibroSoft; PHE, blue; PBS, violet; n=8 PBS, n=15 PHE, ***p=0.0002, Welch’s t test). **D.** Cardiomyocyte cross-sectional area measured via WGA staining. (n=5 visual fields per animal (n=4 PBS, n=7 PHE animals; total of ∼900 cardiomyocytes measured/group, ****p<0.001, Welch’s t test). **E.** Cardiomyocytes analyzed in **D** were further characterized by fibrotic environment (PBS, PHE non fibrotic, and PHE fibrotic) and location of the visual field (longitudinal section: Interventricular septum (IVS), Left Ventricular Wall (LV Wall), and Apex, transverse section: IVS and LV wall; n=4 PBS, n=7 PHE animals, ***p<0.001, ****p<0.0001, analysis done with ImageJ). **F.** Time course assessment of fibrotic deposition and cardiomyocyte cross-sectional area, with end points assessed 24hr post each injection (n=1 PBS, n=3 PHE for each timepoint, WGA measurements of 250 cells/heart section, ****p<0.0001, Welch’s t test).

To investigate the transcriptomes involved in the adaptive response to phenylephrine, we separately isolated myocytes ^20^ and fibroblasts^19^ from cohorts of animals treated as indicated in Figure 1A. Myocytes were enriched to >90% purity as evaluated morphologically^20^ and fibroblasts were ∼87% pure as measured by MEFSK4 positivity by flow cytometry ^19^. RNA sequencing was then conducted on the isolated groups of cells. In cardiomyocytes, hierarchical clustering of the significantly upregulated genes (p_adj_< 0.05, log_2_ fold change ≥ 0.5, total of 1432 genes) followed by Gene Ontology analyses using genomeSidekick ^23^ and gProfiler ^24^ revealed pathways involved in cytoskeletal binding, extracellular matrix (ECM) components, inflammatory responses, and cardiac muscle contraction (Figure 3; note: reported fibroblast-specific genes were excluded, see *Methods*). These findings support the fibrotic phenotype seen by histology, and suggest cardiomyocytes contribute to ECM remodeling as they undergo hypertrophy. Downregulated genes were enriched in pathways including oxidoreductase activity, as well as more general cellular processes like enzyme and signaling receptor binding (Figure 3). The totality of genes up and down regulated in myocytes following phenylephrine treatment are shown in Supplemental Table 1.

**Figure 3.**
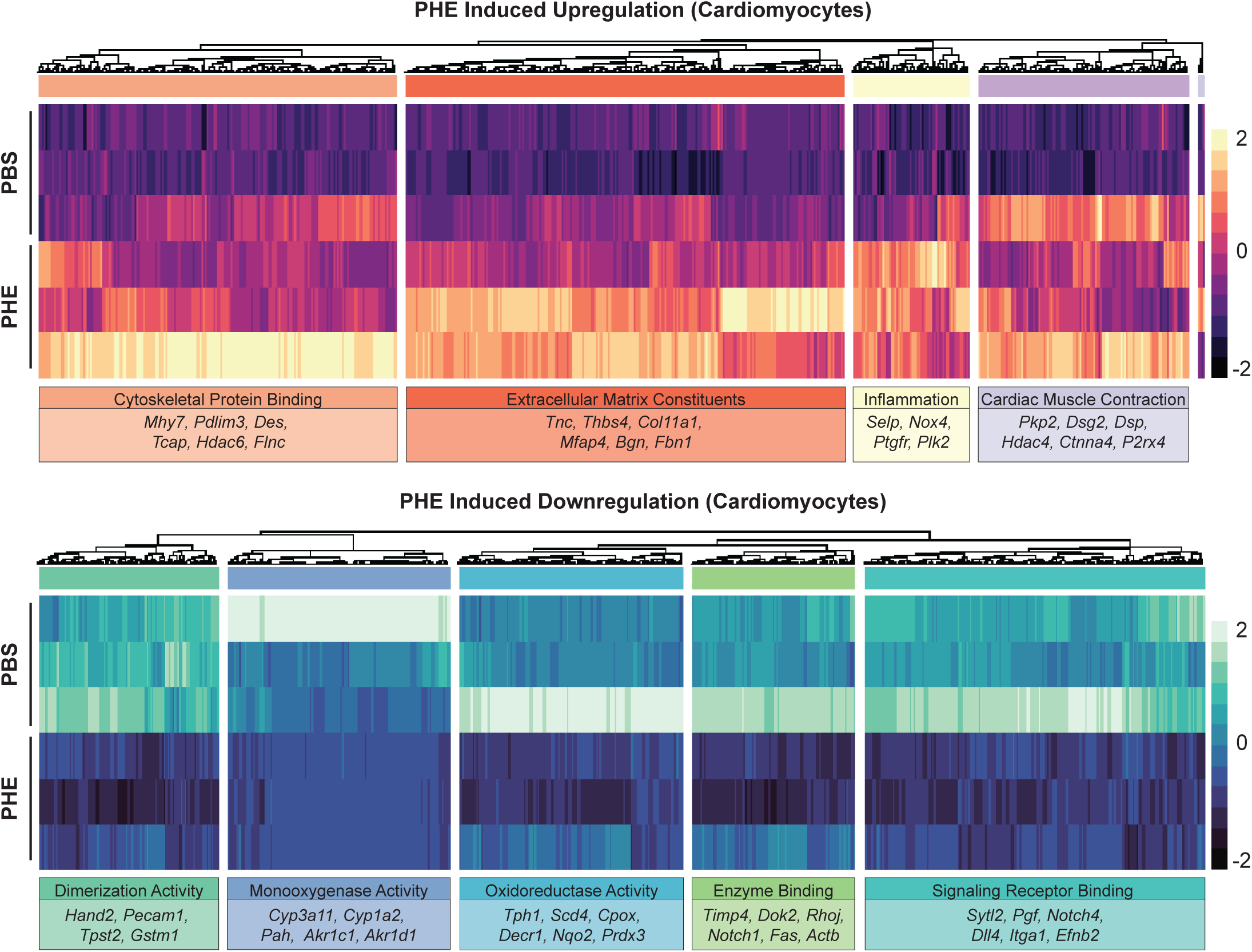
Acute cardiomyocyte transcriptome changes in response to phenylephrine. Top panel shows hierarchically clustered heatmap of significant upregulated genes (p_adj_< 0.05, log_2_ fold change > 0.5, total of 745 genes) subdivided into 4 clusters and top GO terms and representative genes are highlighted. Bottom panel shows heatmap of significant downregulated genes (p_adj_< 0.05, log_2_ fold change < 0.5, total of 893 genes), subdivided into 5 clusters, top GO terms and representative genes highlighted as above.

We next sought to compare the transcriptional signature of acute cardiac stress response to the compensatory phase of heart failure progression as modeled by transverse aortic constriction. Left ventricular pressure overload induced by transverse aortic constriction produces a time-dependent adaptation, characterized by an initial compensatory hypertrophy with little or no change in ejection fraction followed over the course of several weeks by decompensation, as exhibited by further hypertrophy, chamber dilation, wall thinning and depressed ejection fraction ^25, 26^. We investigated overlap between genes altered following 3 days of pressure overload, at which point cardiac hypertrophy is manifest and the hearts not yet in failure as indicated by ejection fraction within normal limits ^25^ and those altered following a week of phenylephrine in this study (Figure 4A). A third of phenylephrine induced gene expression changes were shared with the 3-day pressure overload dataset. Amongst the shared upregulated genes, cytoskeletal proteins, extracellular matrix proteins, and collagen binding proteins were observed, likely indicative of the shared response of the heart to the acute changes in tissue tensile forces and reflecting the accumulation of fibrosis in the hearts following either of these insults (Figure 4A). Intriguingly, the genes upregulated specifically in either the phenylephrine or pressure overload scenario (but not both) clustered into similar gene ontology terms of enzymatic and cytoskeletal protein binding but constituted different individual genes. These findings suggest that phenylephrine induces similar transcriptional remodeling as early pressure overload but often through different pathways (Figure 4B).

**Figure 4.**
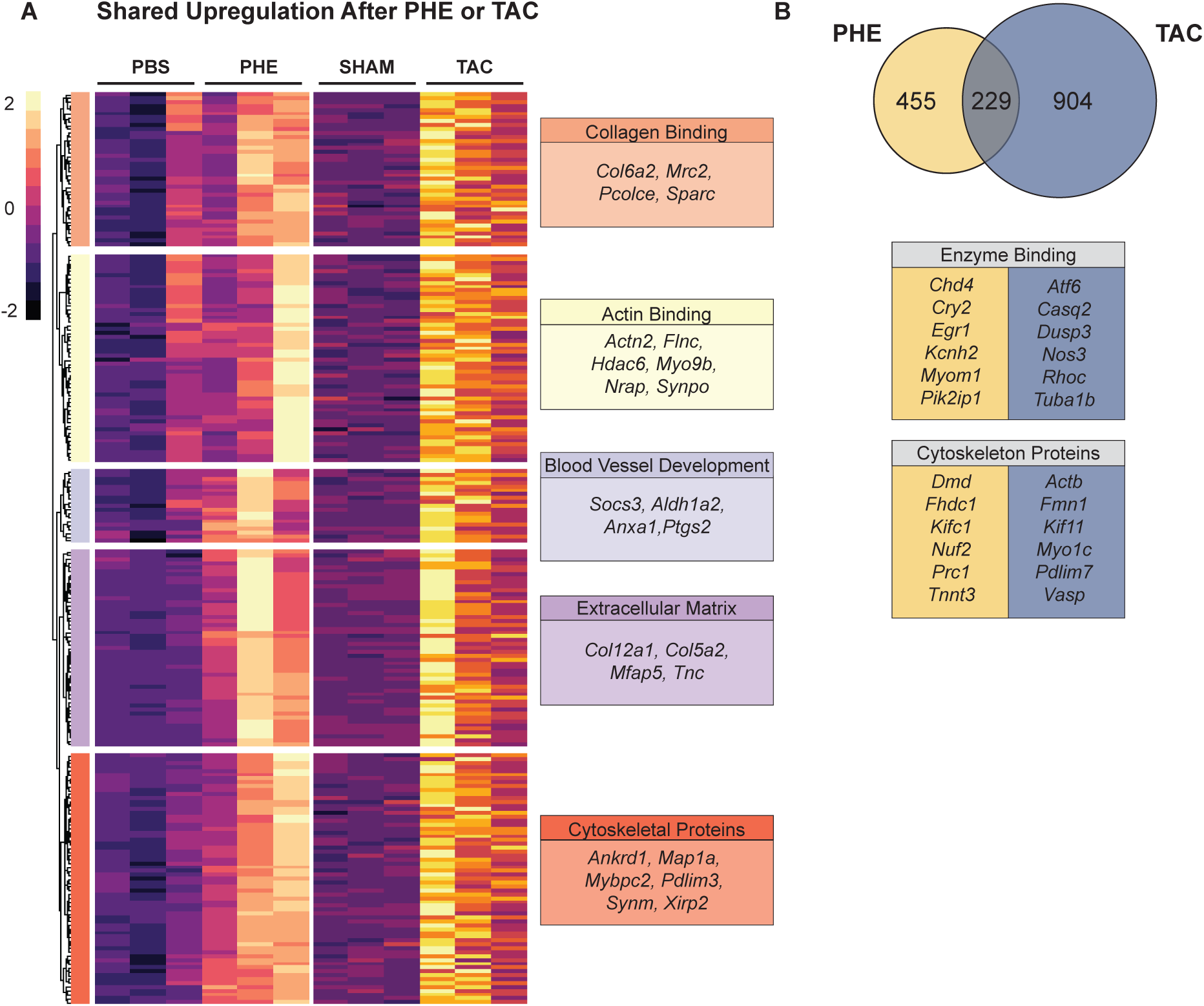
Comparison of cardiomyocyte transcriptional responses between different cardiac stressors. **A.** Upregulated genes in cardiomyocytes following either PHE or transverse aortic constriction (TAC; 3 days). **B.** Venn diagram of all upregulated genes following either PHE or TAC. Note that although the stressors upregulate distinct genes, the ontology terms are shared. Shared upregulated GO terms of genes not found in both PHE and TAC listed in boxes with representative genes from PHE in yellow and TAC in blue. Downregulated genes (shared and distinct) following PHE or TAC are reported in Supplemental Table 1.

We next investigated transcriptome changes in primary fibroblasts isolated from the hearts of phenylephrine treated mice. Upregulated genes were enriched in distinct receptor signaling, extracellular matrix, actin binding, and, unexpectedly, some common myocyte genes (Figure 5), indicative either of myocyte contamination or muscle-like phenotypes in the myofibroblast population ^27, 28^. We reason contamination is unlikely, given that the fibroblast preparations lacked cells resembling morphological myocytes (although it is possible rare atrial myocytes could have been mistaken for fibroblasts). Genes downregulated by phenylephrine in fibroblasts included lipid metabolism, small molecule and protein kinase signaling and a different cohort of cytoskeletal genes (Figure 5). The totality of genes up and down regulated in fibroblasts following phenylephrine treatment are shown in Supplemental Table 2.

**Figure 5.**
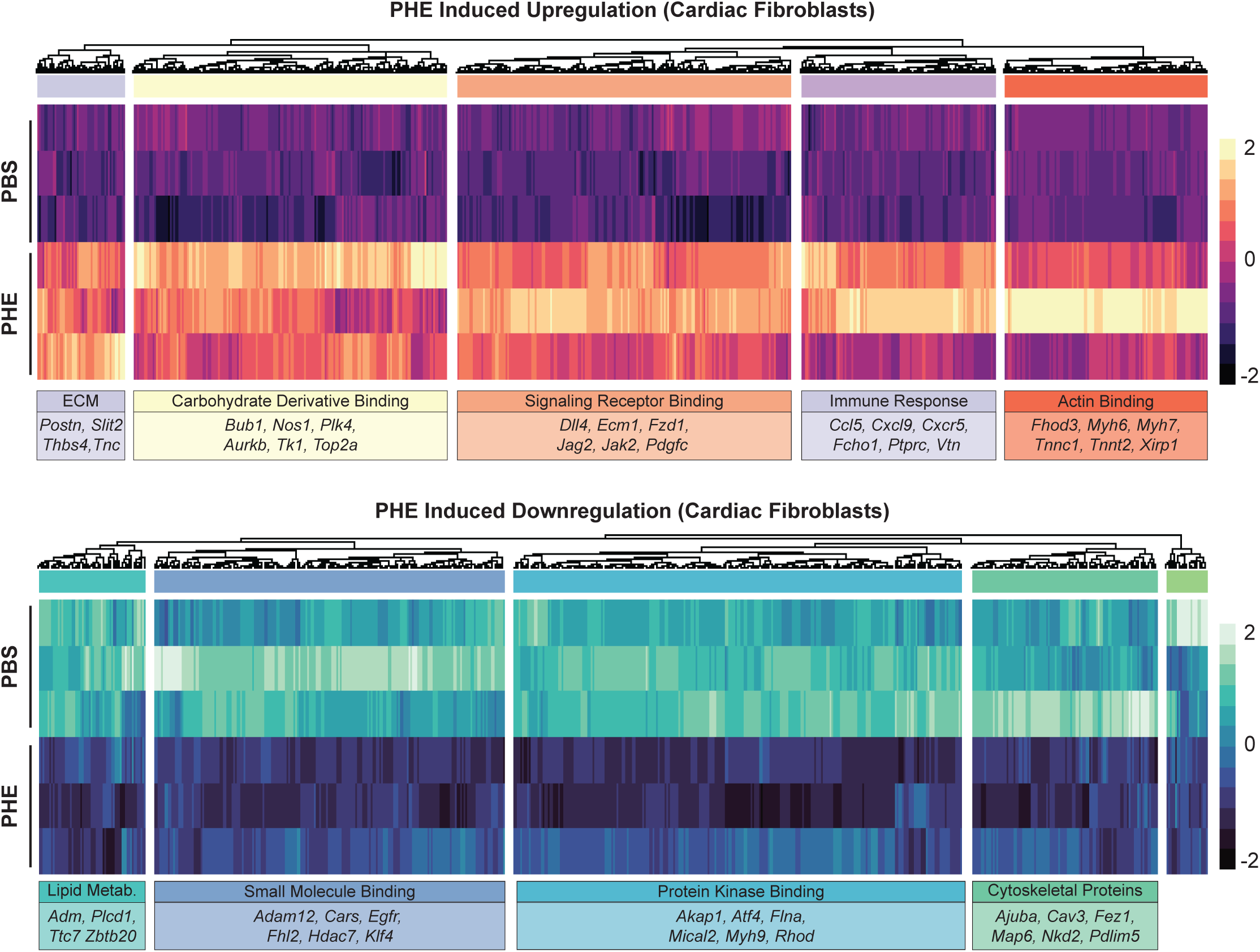
Acute cardiac fibroblast transcriptome changes in response to phenylephrine. Top panel shows upregulated genes (p_adj_< 0.05, log_2_ fold change ≥ 0.5, total of 1589 genes) and bottom panel shows downregulated genes (p_adj_< 0.05, log_2_ fold change ≤ −0.5, total of 550 genes) hierarchically clustered (5 clusters), GO terms and representative genes highlighted on the right.

Next, we sought to investigate overlap in fibroblast gene expression with other models of injury. Upon stress, fibroblasts adopt a myofibroblast phenotype that includes formation of stress fibers, contractile activity and increased secretion of extracellular matrix. This phenotype is observed *in vivo* in heart failure models and can be recapitulated in primary culture by treatment with the cytokine transforming growth factor β (TGF-β). Comparing differentially affected genes between phenylephrine treated hearts with TGF-β treated primary cardiac fibroblasts ^19^ (Figure 6A) showed enrichment in pathways associated with cell adhesion and extracellular matrix production, indicative of the pathophysiological actions of myofibroblasts during stress. However, fewer than 5% of the differentially expressed genes following phenylephrine are shared with the TGF-β treated cohort. Other pathways included intracellular signaling and DNA replication (Figure 6B).

**Figure 6.**
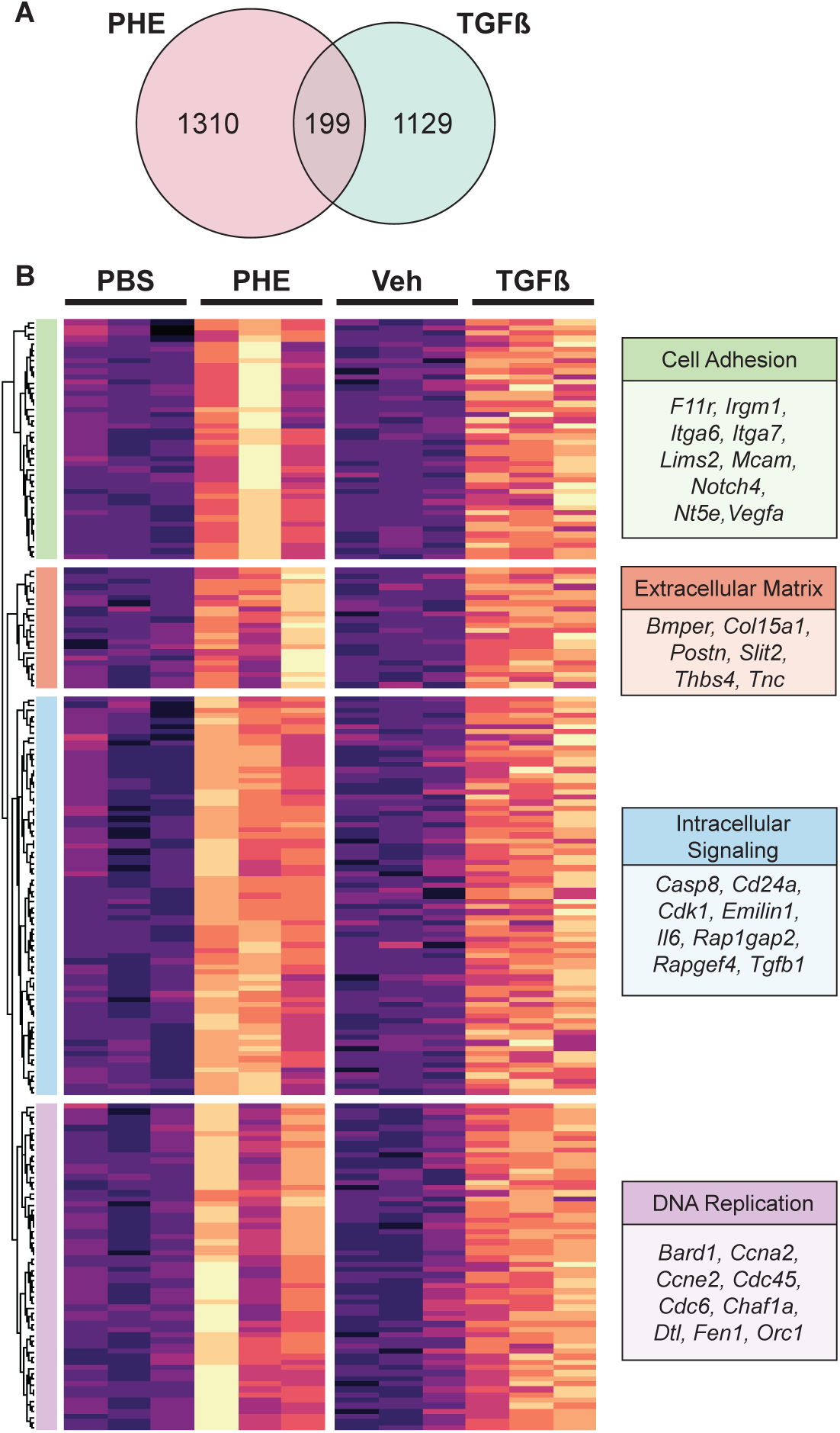
Comparison of cardiac fibroblast stress responses between different stressors. **A.** RNA-seq cardiac fibroblasts from PHE treated mice were compared to TGFβ treated fibroblasts. **B.** The shared upregulated genes were hierarchically clustered (4 clusters) and GO terms on those clusters revealed a subset highlighted by DNA replication.

We next investigated the transcription factors likely to be remodeling the cardiac transcriptome after phenylephrine treatment. We used HOMER to determine enriched motifs in the proximal promoter regions of the up-regulated genes from phenylephrine treated hearts. In myocytes, TEAD3, GLIS3, RUNX1/2, PGR, FOS and HLF were some of the top enriched motifs, whereas phenylephrine-induced cardiac fibroblast genes were enriched for motifs from the IRF, ETS, and MEF2 transcription factor families (Figure 7).

**Figure 7.**
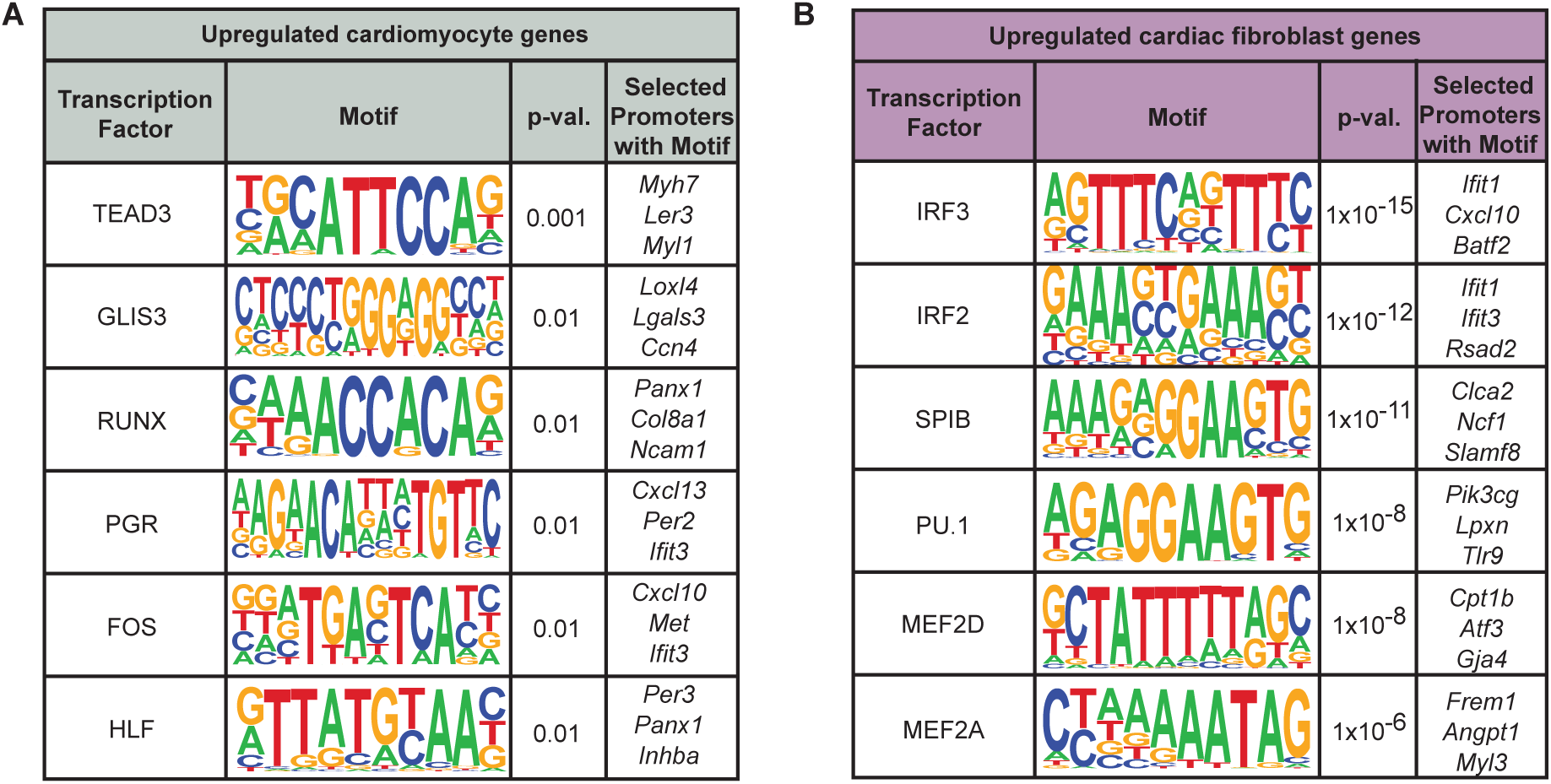
Transcription factor motif analysis of genes upregulated in phenylephrine-treated hearts. **A.** Motif analysis of the promoter regions of genes significantly upregulated (p_adj_ < 0.05 and log_2_(fold change) > 1) in cardiomyocytes isolated from phenylephrine-treated hearts compared to PBS controls. **B.** Motif analysis of the promoter regions of genes significantly upregulated (p_adj_ < 0.05 and log_2_(fold change) > 1) in cardiac fibroblasts isolated from phenylephrine-treated hearts compared to PBS controls.

To investigate a potential transcriptional relationship corresponding to the cellular cross talk between the extracellular matrix deposition and hypertrophy, we employed the computational platform NicheNet ^29^, which uses experimentally determined protein interaction pathway data to predict communication between cells on the basis of a ligand’s expression in one cell and its targets’ activation in another (Figure 8A). To examine the fibroblast to cardiomyocyte signaling pathway, we began by generating a list of prioritized fibroblast ligands that are predicted to significantly upregulate cardiomyocyte genes that we observed in our RNA-seq dataset (Figure 8B). TGF-β was one of the top ligands and was also upregulated in our fibroblast RNA-seq dataset. Next, we graphed the known regulatory networks downstream of TGF-β and measured the level of expression of ECM target genes in our respective datasets, simultaneously capturing both the strength of the regulatory interaction and involvement of signaling intermediates in the target cells (Supplemental Figure 2).

**Figure 8.**
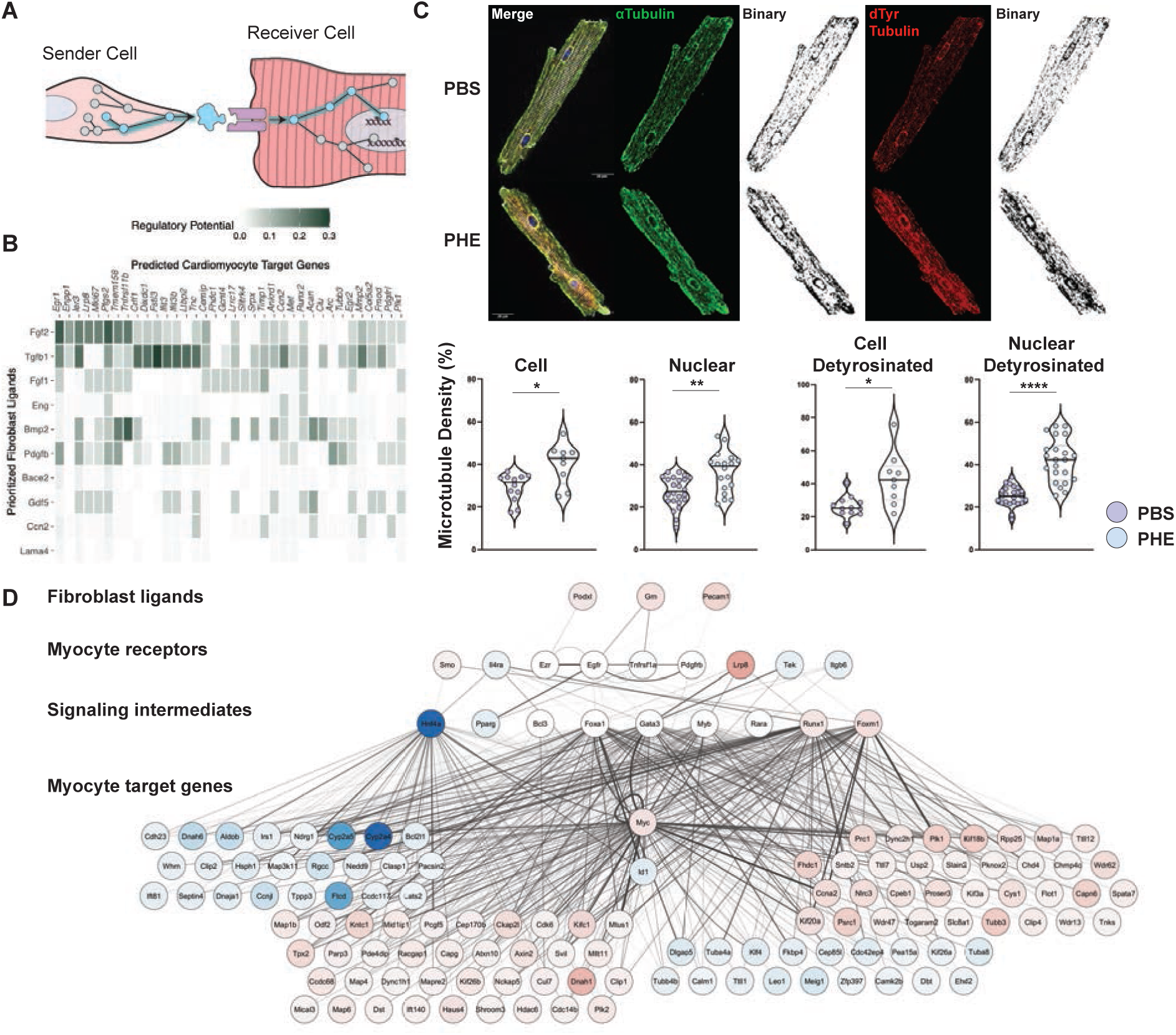
Cardiomyocyte and fibroblast cross talk in phenylephrine treated hearts. **A.** Schematic of cardiac fibroblast-to-myocyte communication analysis. **B.** Prioritized fibroblast-derived ligands and their predicted gene regulatory targets in cardiomyocytes. Heatmaps are colored by the weight of the ligand-target regulatory potential in the NicheNet prior model. **C.** *Top,* Confocal z-stack (4 µm merge around the nuclear planes) imaging of the microtubule network in PBS (top) and PHE (bottom) isolated cardiomyocytes (scale = 20 micron). *Bottom,* α-tubulin density compared to total cell area, PHE (blue) compared to PBS (violet). Detyrosinated tubulin quantification performed in analogous manner (n=5 visual fields per cell; n=2 PBS, n=2 PHE animals, total of 10 cardiomyocytes measured/group, *p=0.01, **p=0.001, Welch’s t test). **D.** Ligand-receptor analysis of microtubule-related changes in cardiomyocyte gene expression. Nodes are colored by change in transcript abundance in fibroblasts (for ligands) or cardiomyocytes (for receptors, signaling intermediates, and target genes) from phenylephrine-treated hearts. Edges are weighted by strength of the signaling or gene regulatory interaction the NicheNet prior model.

To investigate the mechanism of cardiomyocyte hypertrophy in response to phenylephrine, we examined microtubule dynamics. Cardiomyocyte passive stiffness is largely due to the sarcomeric spanning protein titin ^30^. The microtubule cytoskeleton plays a role in providing tension to the sarcomeric contraction-relaxation cycle and microtubules are stabilized and more resistant to compression in heart failure patients ^30^. Cardiomyocytes from phenylephrine or PBS treated animals were stained for α-tubulin or detyrosinated tubulin—a post-translationally modified tubulin that binds to the sarcomeric z-disk. Detyrosinated tubulin is a more stable species of the protein and is upregulated in congenital hypertrophic cardiomyopathy, resulting in resistance of the muscle to compression and relaxation ^30^. Confocal imaging was performed, merging 4 µm regions around the nuclear plane, thresholding for a binary image and assessing microtubule density compared to total cell area (Figure 8C). Both α-tubulin and detyrosinated tubulin were modestly dense throughout the whole cell in agreement with previous studies ^18^, but interestingly, localized measurements around the nuclear area revealed the deposition of α-tubulin and detyrosinated tubulin to be significantly denser, suggesting a local network of microtubules around the nucleus creates a localized stiff nuclear environment. To explore the intercellular communication underpinning this phenotype, we generated a predicted network of microtubule regulation based on protein interaction data. Because fibrosis preceded hypertrophy at the tissue level, we hypothesized that the growth signals to myocytes emanate from fibroblasts. The resulting network revealed that genes for many of the known protein interactors in the microtubule network underwent altered expression following phenylephrine, while several known constituents of this network were unchanged (Figure 8D). Many transcription factors identified in our motif analysis (Figure 7) appear as likely signaling intermediates in this network. This analysis enables integration of prior knowledge regarding specialized protein interaction networks with new primary data, revealing the specific circuits operative in response to distinct stressors.

## Discussion

Myocyte hypertrophy and cardiac fibrosis are central features of heart failure. These phenotypes have many possible drivers in the setting of heart failure including myocyte death, local and systemic inflammation driven by cytokines and immune cells, and berserk autonomic signaling. In some subjects, infarction is a major contributor to the initial injury, usually leading to heart failure with reduced ejection fraction (HFrEF), whereas in the other half of subjects whose ejection fraction is preserved (HFpEF), hypertension and metabolic syndrome are commonly the precipitating causes. Both HFrEF and HFpEF are accompanied by cardiac fibrosis ^3^, although given the complexity of the pathology in humans, the relative contribution of various fibrotic stimuli to this phenotype, as well as the actions of different cells in the heart, will vary between individuals. To address this issue, we sought to examine the role of aberrant adrenergic stimulation—a hallmark of both HFrEF and HFpEF—on cardiac function, structure, tissue remodeling and cell type-specific transcription.

In this manuscript we show that upon acute cardiac hypertrophic stress with phenylephrine, fibrosis occurs first, followed by myocyte hypertrophy, suggesting extensive intercellular crosstalk that we highlight through cell-to-cell interactome analyses. Acute phenylephrine administration induced a rapid development of fibrosis, peaking one day after treatment and not expanding over the ensuing 7 days. Myocyte hypertrophy was slightly delayed, peaking at 3 days, demonstrating a time course similar to that reported in other models of cardiac pathology *in vivo* (e.g. pressure overload ^31^) and following phenylephrine treatment of myocytes in culture ^32^. Combined histological analyses indicated that in addition to a global increase in fibrotic area, the regions of myocyte hypertrophy were co-localized with regions of fibrosis within the ventricle, demonstrating an anatomical link between these distinct biomechanical mechanisms of compensation.

Our model does not induce heart failure, but instead simulates early transcriptional remodeling after acute stress. We observe pathologic features shared by both HFrEF and HFpEF including fibrosis, hypertrophy, and diastolic dysfunction, suggesting that α-adrenergic activation may be an underlying factor in these aspects of heart failure. Phenylephrine administration and enhanced α-adrenergic stimulation also decreases capillary density in the heart ^5^, which may also be factors contributing to the phenotypes we observed in this study. Because our model occurs acutely and in the absence of sustained hypertension, α-adrenergic receptor antagonists may be a possible mechanism to modulate fibrosis in patients with dysregulated autonomic control. Recent work with a similar model of acute phenylephrine administration has identified the cytoskeletal rearrangements required for precise transcription and translation of proteins necessary for myocyte hypertrophy ^18^. Our study adds to these and other observations in response to adrenergic simulation (such as those examining isoproterenol and phenylephrine in concert^33^), by characterizing the tissue ultrastructural remodeling during acute α-adrenergic stimulation and determining the totality of transcriptional reprogramming underpinning these phenotypes.

Intercellular communication is a hallmark of fibrosis. Cytokine signaling between myocytes, immune cells, fibroblast and other populations drives ECM remodeling at various stages of heart failure ^34^, which we also observed in response to phenylephrine in our cell-cell communication analyses. While fibroblasts undoubtedly play a central role in fibrosis and ECM production, determining the source of ECM proteins themselves remains an open area of investigation. Depletion of fibroblasts from the mouse heart (using a PDGFRα-driven diphtheria toxin A strategy) revealed these cells to be dispensable for normal cardiac function: hearts virtually devoid of fibroblasts in the PDGFRα lineage functioned normally and had minimally modified ECM, as measured by hydroxyproline quantification, electron microscopic analysis, and ECM proteomics^35^. This surprising finding suggests that cells other than fibroblasts can contribute to the maintenance of ECM in the healthy adult mouse heart. In the same study, the authors showed that adult mice lacking PDGFRα+ fibroblasts exhibited impaired collagen production and enhanced myocyte hypertrophy following infusion of angiotensin II and phenylephrine ^35^, further highlighting the intercellular communication between myocytes and fibroblasts in response to pathologic stress and implying that this communication is essential for fibrosis (and that, perhaps in the absence of fibrosis, myocytes undergo additional hypertrophic growth in compensation).

Our findings add to the understanding of this myocyte-fibroblast cross talk at the level of *in vivo* phenotypes and provide curated transcriptomes to allow the community to interrogate the cell type-specific changes in gene expression, as well as the intercellular crosstalk, as demonstrated by our ligand-receptor analyses. While single cell or single nucleus RNA sequencing approaches are valuable, these studies inherently lack orthogonal validation of the cell types in question. Rather, cell type assignment following dimension reduction analyses of single cell or single nucleus data proceeds based on the experimenters’ knowledge of the cell types: that is, marker genes are selected based on prior literature and used to annotate cell clusters ^7^. Our approach is complementary, in that we started by purifying myocyte or fibroblasts from the other cells in the heart and separately analyzed them. This approach revealed an unexpectedly large contribution of myocytes to the production of ECM and enabled a comparison of the phenylephrine-responsive transcriptome with that mobilized following either pressure overload or direct stimulation of fibroblasts with TGF-β. This study will enable an improved understanding of the phenotypic progression and intercellular cross talk in response to the various individual stimuli that induce cardiac dysfunction and contribute to heart failure.

## Methods

### Phenylephrine Treatment

All animal surgery, echocardiography and euthanasia procedures were approved by the UCLA Animal Research Committee in compliance with the National Institutes of Health Guide for the Care and Use of Laboratory Animals. C57Bl/6J male mice adult (8-10 weeks old) were used in the study. Phenylephrine was administered (20 mg/kg in PBS, approximate total volume 100ul, depending on body weight) on day 1 and every other day until day 7 by subcutaneous injection.

### Echocardiography

Heart function was measured by echocardiology at basal and on day 7 of phenylephrine treatment. Animals were anesthetized with 1.5% isoflurane and 95% O2, and chest hair was removed. Heart rates were maintained between 400 and 500 beats per minute using continuous ECG monitoring. Body temperature was set at 37 °C using a heating pad. The Vevo 3100 (VisualSonics) was used to collect left ventricular (LV) systolic measurements included long and short axis B-mode imaging and short axis M-mode for ejection fraction, LV wall thickness and chamber diameter. Diastolic measurements for the LV were taken from the 4-chamber apical view, with E/A ratio from the pulse wave doppler and E/e’ from tissue doppler. All calculations were performed using the Vevo LAB 5.6.1software.

### Invasive Hemodynamics

Hemodynamics measurements were performed on phenylephrine (n=5) and PBS (n=5) mice. Mice were intubated and anesthetized with 2% inhaled isoflurane. Three ECG leads were used to obtain heart rate and RR interval recordings. A midline incision was made in the neck to expose the right carotid artery. A 1F Millar catheter (Millar Medical) was inserted into the right carotid artery and advanced until an arterial pressure waveform was obtained for continuous blood pressure recordings. The catheter was then advanced into the LV chamber for measurement of left ventricular end-diastolic pressure (LVEDP). After hemodynamic assessment the isoflurane was increased to 5% and the animals underwent cardiectomy with anesthesia.

### Tissue Section Histology

Cardiac tissue samples were fixed in 10% formalin buffered solution (Sigma: HT501128) overnight, dehydrated in 70% ethanol and sent to the UCLA Translational Pathology Core to generate paraffin blocks. Samples were cut into 4 µm thick slices, put on slides, and stained with hematoxylin and eosin, Masson’s trichrome stain (Sigma: HT15-1KT), and Picro Sirius (abcam: ab150681) to detect fibrosis. Fibrotic area for each slide was quantified and expressed as the percentage of the area occupied by the whole heart on a given slide using FibroSoft ^36^. For cardiomyocyte cross-sectional area, cell size was determined using Zeiss Axio Vert.A1 on heart slices labeled with wheat germ agglutinin (WGA) (Thermo: W11262). Images were taken with the 20x objective, taking 5 image fields from each tissue area (LV septum, LV wall, and apex (4-chamber view only)) and cell size was traced using ImageJ. To further characterize cardiomyocyte size in relation to fibrotic area, each image was scored as fibrotic or non-fibrotic based on WGA morphology. Tracings were separated and plotted using GraphPad.

### Cardiomyocyte Isolation

Using an established protocol ^20^, adult mice were treated with heparin (100 USP units) for 20 min to prevent blood coagulation followed by anesthetization with sodium pentobarbital (100 µl of 50 mg/ml dilution, intraperitoneal). Upon loss of rear foot reflex, the heart was removed and instantaneously arrested in ice-cold phosphate buffered saline (PBS). The aorta was canulated with a blunt ended 22-gauge needle and mounted on a modified Langendorff apparatus for perfusion with Tyrode’s solution, digestion buffer, and Krebs buffer (KB). Cardiomyocytes were dissociated in KB solution, filtered (100 µm strainer, 25 µM blebbistatin added if used for imaging purposes) and centrifuged 2 min at 200xg for further usage. This method obtained cells that were ≥95% cardiomyocytes by visual inspection of rod-shaped cell morphology.

### Fibroblast Isolation

Primary cardiac fibroblasts were isolated using enzymatic digestion (7 mg/ml collagenase type II: Worthington Biochemical Corporation: LS004177) followed by centrifugation (2000 rpm for 8 min at 4°C) and cell plating in DMEM/F12 media supplemented with 10% fetal bovine serum (FBS), 1% antibiotics (penicillin and streptomycin), and 0.1% insulin-transferrin-selenium (ITS; Corning: 354350). After 2 h, cells were maintained in DMEM/F12 media supplemented with 10% fetal bovine serum (FBS), 1% antibiotics (penicillin and streptomycin), human basic fibroblast growth factor (hbFGF, 1:10000 concentration from 200X stock; Millipore Sigma: 11123149001) and 0.1% insulin-transferrin-selenium (ITS; Corning: 354350). Cells were fixed 18 h later for confocal imaging.

### RNA-seq and Data Analyses

Left ventricular cardiomyocytes (PBS: n=3, PHE: n=3) and fibroblasts (PBS: n=3, PHE: n=3) were isolated and flash frozen for RNA-sequencing. The Technology Center for Genomics and Bioinformatics Core (UCLA) isolated RNA from the cells. RNA-seq library preparation was performed by the Technology Center for Genomics & Bioinformatics Core (UCLA) following standard protocols including removing ribosomal RNA using an Illumina Ribo-Zero rRNA Removal Kit. Libraries were sequenced in an Illumina NovaSeq 6000 at a minimum of 40 million reads per sample.

Data analysis was performed using custom pipelines. Paired-end 150 bp reads were demultiplexed into sample-specific *_R1.fastq and *_R2.fastq files using custom scripts and then pseudoaligned to the mm10 genome using the ultra-fast pseudoalignment tool, Salmon ^37^. Briefly, we ran salmon index to generate a mm10 (mouse) transcriptome index using the appropriate “known cDNA” FASTA file from Ensembl release 8117. Salmon quant with mm10-specific index was used to pseudoalign corresponding read pairs to the appropriate transcriptome, using *_R1.fastq and *_R2.fastq files as inputs. This method generates a transcript count for each known gene in the genome to be used in downstream calculation of count-based statistics.

Salmon transcript quantifications (*.sf files) for all samples were loaded into R (version 4.3.2) using tximport, filtering out genes with 0 detected transcripts. The data from each cell type was summarized into a separate DESeqDataSet using DESeq2’s ^38^ DESeqDataSetFromTximport() function. Each DESeqDataSet was filtered to exclude genes with fewer than 10 transcripts across conditions before differential gene expression was performed using DESeq2’s DESeq() function, with default parameters. A data frame containing normalized count data and the DESeq results was created for downstream analyses.

### Gene Ontology Analysis

Gene Ontology analysis was performed on the significantly upregulated (p.adj.<0.05 & log2FC>1) genes between phenylephrine and PBS conditions using g:Profiler’s ^24^ gost() function with the mmusculus organism and gSCS correction method. Before running the cardiomyocyte dataset, we removed 15 genes (*Ccna2, Col1a1, Col1a2, Col3a1, Col5a1, Ereg, Fn1, Lum, Myc, Nlrc3, Pak3, Postn, Ptprv, Serpine1,* and *Sfrp1*) intersecting with Supplemental Table 3, the genes collated from a literature search and Gene Ontology terms primarily associated with cardiac fibroblast activation (GO:0072537), migration (GO:0010761), and proliferation (GO:0048144).

### Motif Analysis

Motif analysis of the promoters of the significantly upregulated (p.adj.<0.05 & log2FC>1) cardiomyocyte and cardiac fibroblast genes was conducted with HOMER’s findMotifs.pl function, using the entire mouse reference promoter set as background. Before running the cardiomyocyte dataset, we removed 15 genes (*Ccna2, Col1a1, Col1a2, Col3a1, Col5a1, Ereg, Fn1, Lum, Myc, Nlrc3, Pak3, Postn, Ptprv, Serpine1,* and *Sfrp1*) intersecting with Supplemental Table 3, a table of genes collated from a literature search and Gene Ontology terms primarily associated with cardiac fibroblast activation (GO:0072537), migration (GO:0010761), and proliferation (GO:0048144). To find promoters that contain the significantly enriched motifs, we ran findMotifs.pl again with the - find parameter and the knownMotif.motif file of each motif of interest. Highlighted genes at promoters of interest in Figure 7 were chosen by motif score and topic relevance.

### Cell-Cell Communication Analysis

NicheNet ^29^ was used for cell-cell communication analysis. NicheNet’s mouse ligand-receptor network, ligand-target matrix, and weighted networks were read into R v4.3.2, along with the dataframes containing cardiac fibroblast and myocyte normalized RNA-seq counts and the DESeq2 results. For cardiomyocyte to fibroblast analyses, cardiomyocytes were designated the “sender” cells and fibroblasts the “receiver” cells, and vice versa for fibroblast to cardiomyocyte analyses. For both analyses, a set of potential ligands were identified from the sender cell type by intersecting the ligands from the ligand-target matrix with the genes expressed (p.adj. not NA) in receiver cells. These potential ligands were filtered to only include those that bind a receptor in the ligand-receptor network that is expressed in the receiver cell type (p.adj. not NA). Targets in the receiver cell type were defined as genes significantly upregulated in phenylephrine-treated hearts (p.adj.<0.05 & log2FC>1) that appear in the mouse ligand-target matrix. Cardiomyocyte targets were further filtered to exclude the 15 myofibroblast-associated genes that intersected with Supplemental Table 3, (*Ccna2, Col1a1, Col1a2, Col3a1, Col5a1, Ereg, Fn1, Lum, Myc, Nlrc3, Pak3, Postn, Ptprv, Serpine1,* and *Sfrp1*). Ligands regulating the expression of the target genes in the receiver cell type were predicted with NicheNet’s predict_ligand_activities() function, which ranks ligands based on how well they predict target gene expression. Ligands were ranked by Pearson correlation coefficient and the top 10 (“prioritized ligands”) were used for downstream analysis. Receptors of each prioritized ligand were retrieved from the ligand-receptor network. The top 100 targets of each prioritized ligand were retrieved from the ligand-target matrix and merged with differential gene expression data to create Supplemental Tables 4 and 5. To visualize the signaling and gene regulatory network connections involved in inflammatory pathways in cardiac myocytes and fibroblasts in Figure 8, we intersected the Gene Ontology term GO:0010467 (GO:BP Cytokine Production) with cardiomyocyte targets and GO:0002376 (GO:BP Immune System Process) with fibroblast targets to create a gene set of inflammation-related targets for each receiver cell type. We used these gene sets of interest, along with the significantly upregulated ligands (p.adj.<0.05 & log2FC>0) from the sender cell type’s prioritized ligands, as the input for NicheNet’s get_ligand_signaling_path() function, considering the intersection between the top 2 targets of a ligand and the gene set of interest. This network was loaded into Cytoscape ^39^ and the width and opacity of the networks’ edges were weighted by the weight of the interaction. A node annotation table was also created and uploaded to Cytoscape to assign each gene a role as a ligand, target, receptor (lr_network$to), or signaling intermediate (all other nodes in signaling or regulatory outputs of get_ligand_signaling_path()), and was used for spatial grouping of the nodes in Cytoscape. Similarly, a node expression table was created and uploaded to Cytoscape to color nodes by significant up- or down-regulation in phenylephrine-treated hearts compared to PBS in receiver cells (if the node was annotated as a receptor, signaling intermediate, or target) or sender cells (if the node was a ligand).

### Quantification and Statistical Analysis

Data presented as the mean +SD, unless otherwise indicated in the figure legends. Statistical analyses were performed using Prism software v9.0 (GraphPad Software) using Welch’s t-test between two groups and one-way ANOVA with Tukey’s multiple comparison analysis between three or more groups. A p-value less than 0.05 was considered statistically significant.

## Data Availability Statement

RNA-seq data used in this manuscript will be available upon publication. Note: Please email Thomas Vondriska for Supplemental Tables.

## Acknowledgements

We thank the Technology Center for Genomics and Bioinformatics Core for RNA sequencing experiments and the Pathology Core for immunohistochemistry specimen preparation.

## Author Contributions

THK, TG, KW: animal physiology and analysis of cardiac function; THK, TG, KW, MV, MAF, DJL, TMV: physiology data analysis and interpretation; THK, NDG, TG, DJC: RNA-seq data analyses; THK: microscopy; DJL, TMV: infrastructure and funding; THK, DJL, MAF, TMV: inception, study design and interpretation; THK, TG, NDG, TMV: figures; TMV: writing; all authors edited and approved the final version.

---

**Supplemental Figure 1.**
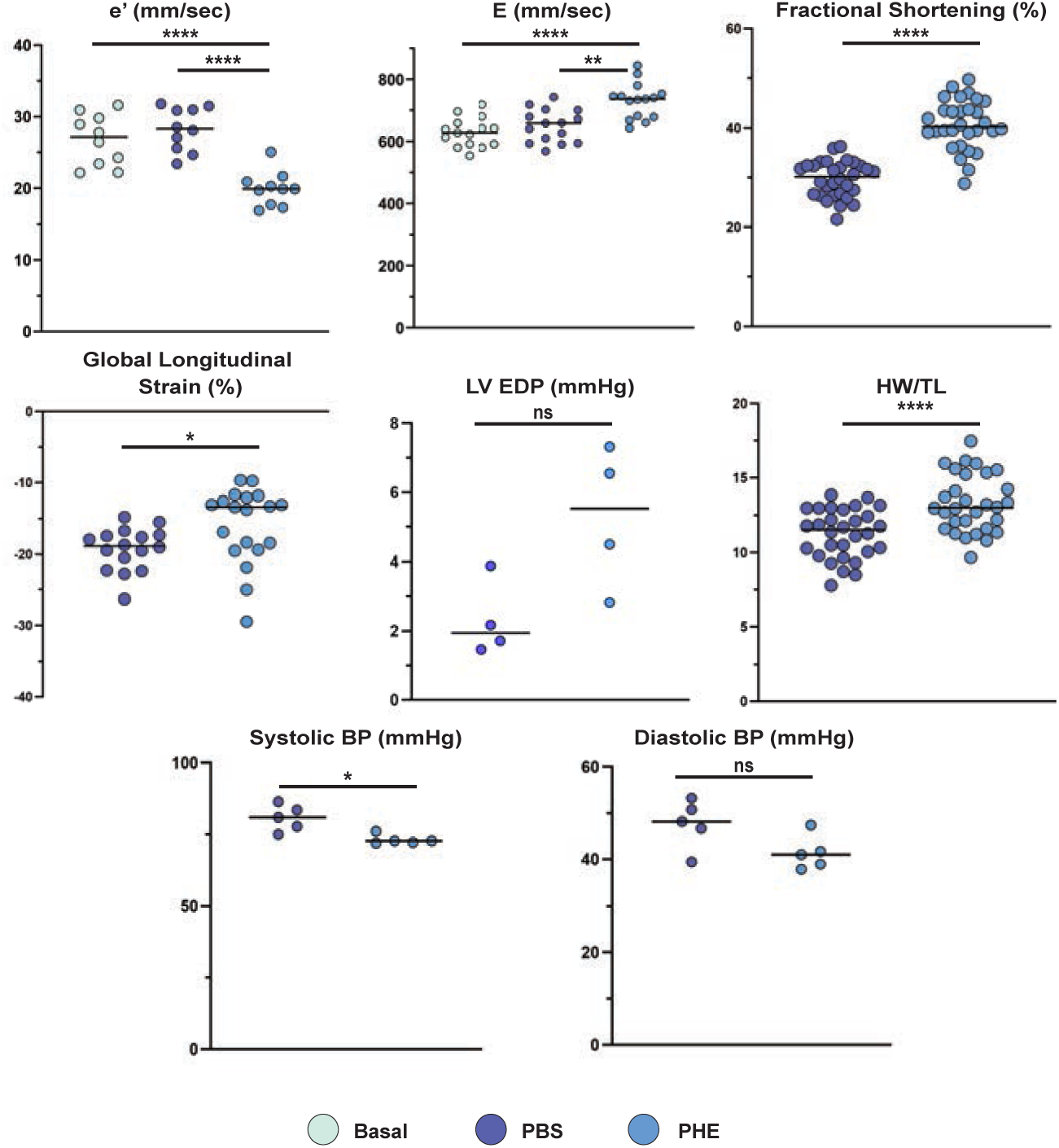
Cardiac function and structure measurements in response to phenylephrine. PHE (blue), basal (green) and 1 week PBS (purple). For e’, E, and FS: n= 30 mice/group, ****p<0.0001, One Way ANOVA. For global longitudinal strain (GLS): n = 16 PBS, n=20 PHE, *p<0.0001, Welch’s t test. For heart weight (mg):tibia length ratio (mm) (HW/TL): n= 30 mice/group, ****p<0.0001, Welch’s t test. For invasive hemodynamics including LV end diastolic pressure (LV EDP), diastolic blood pressure and systolic blood pressure: n=5, *p<0.05, Welch’s t test.

**Supplemental Figure 2.**
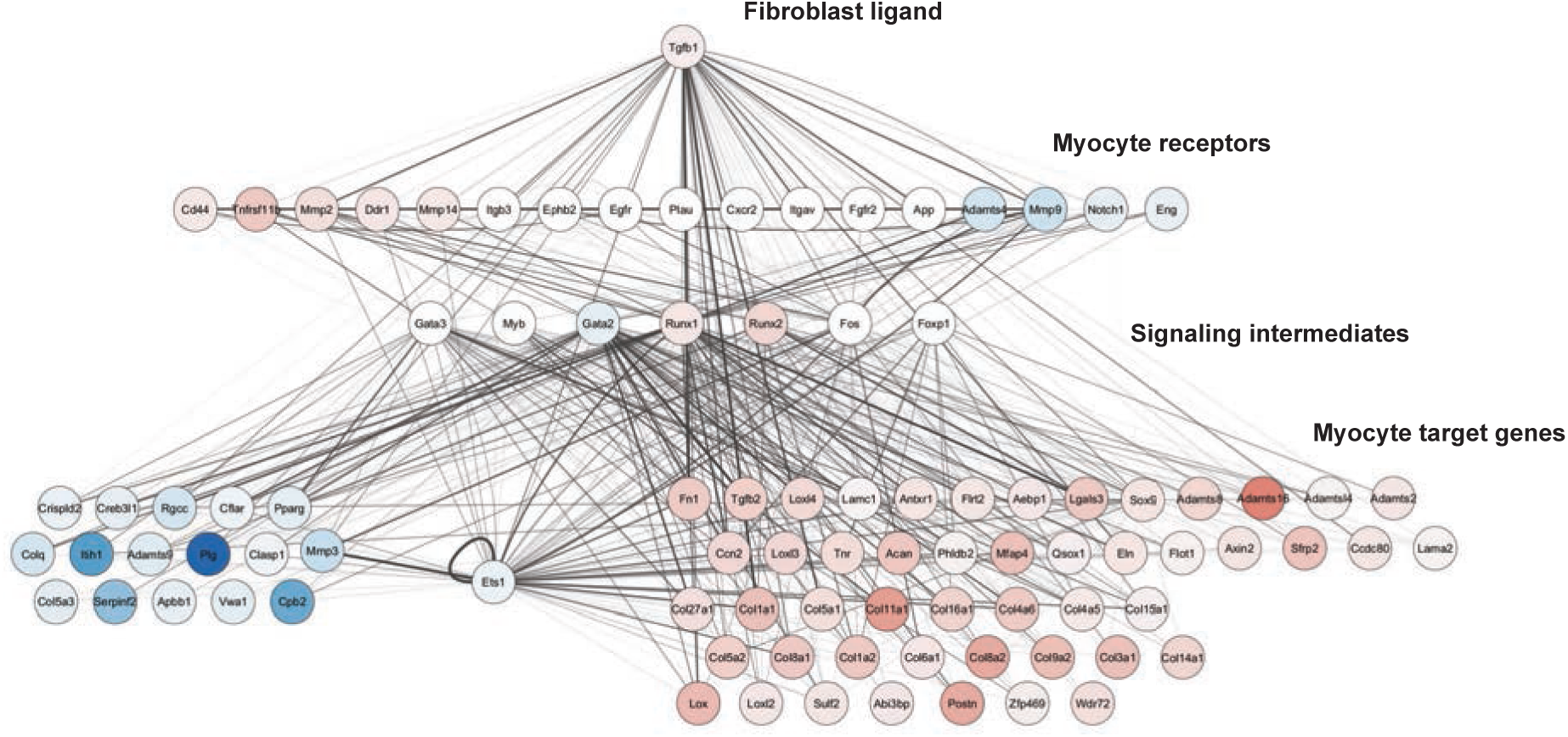
Myocytes contribute to ECM production and organization in response to phenylephrine. Ligand-receptor analysis of the ECM-organization-related gene expression changes with phenylephrine in cardiac myocytes highlight a role for TGFβ signaling in this response. Nodes are colored by change in transcript abundance in fibroblasts (for ligands) or cardiomyocytes (for receptors, signaling intermediates, and target genes) from phenylephrine-treated hearts. Edges are weighted by strength of the signaling or gene regulatory interaction the NicheNet prior model.

